# Coordinated emergence of hippocampal replay and theta sequences during post-natal development

**DOI:** 10.1101/505636

**Authors:** Laurenz Muessig, Michal Lasek, Isabella Varsavsky, Francesca Cacucci, Thomas J. Wills

## Abstract

Hippocampal place cells encode an animal’s current position in space during exploration[1]. During subsequent sleep, hippocampal network activity recapitulates patterns observed during recent experience: place cells with overlapping spatial firing fields during locomotion show a greater tendency to co-fire (‘reactivation’) [2] and temporally ordered and compressed sequences of place cell firing observed during wakefulness are reinstated (‘replay’) [3–5]. Reactivation and replay are thought to be network mechanisms underlying memory consolidation [6–10].

---

Compressed sequences of place cell firing also occur during exploration: during each cycle of the theta oscillation, the set of active place cells shifts from those signalling positions behind to those signalling positions ahead of an animal’s current location [11,12]. These ‘theta sequences’ have been linked to spatial planning [13].

Here we demonstrate that, before weaning (post-natal day 21, P21), offline place cell activity reflects predominantly stationary locations in recently visited environments. By contrast, sequential place cell firing, describing extended trajectories through space during exploration (‘theta sequences’) and subsequent sleep (‘replay’), emerge gradually after weaning in a coordinated fashion, possibly due to a protracted decrease in the threshold for experience-driven plasticity.

Hippocampus-dependent learning and memory emerge late in altricial mammals [14–17], appearing around weaning in rats and slowly maturing thereafter [15]. In contrast, spatially localised firing can be observed at least one week earlier (albeit with reduced spatial tuning/stability) [18–21]. By examining the emergence of hippocampal reactivation, replay, and theta sequences during development, we show that the coordinated maturation of offline consolidation and online sequence generation parallels the late emergence of hippocampal memory in the rat.

## RESULTS

We first investigated the development of reactivation, defined as changes in cell-pair firing correlations following exploration (Figure 1a, see Methods). We recorded 1566 complex spike (CS) cells from region CA1 from 24 animals aged between P17 and P32 as they ran in a familiar square open field environment (RUN) and during the sleep phase immediately preceding (PRE-sleep) and following (POST-sleep) exploration, yielding a total of 19,334 cell pairs. From P17 onwards, the similarity of the place fields of CS cell pairs during RUN was significantly correlated with their co-activity during Sharp-Wave Ripples (SWRs; see Methods), selectively during POST-sleep (but not during PRE-sleep; Figure 1b, c). The activity of hippocampal principal neurons that fire together during the exploratory phase is therefore selectively reinstated during sleep periods directly following exploration in young rats, similarly to what is observed in adult rodents [2]. Indeed, at all ages, RUN versus sleep co-firing correlation was significantly increased in POST– vs PRE-sleep (Figure 1c). Similar results are obtained when the similarity of cell pair RUN co-firing is assessed at the finer time scale of single theta cycles (Figure 1d). These results demonstrate that Hebbian plasticity between hippocampal CS cell pairs is present from the earliest ages tested in the rat: neurons that fire together during RUN show an increased propensity to closely timed co-firing during post-experience sleep.

**Figure 1.**
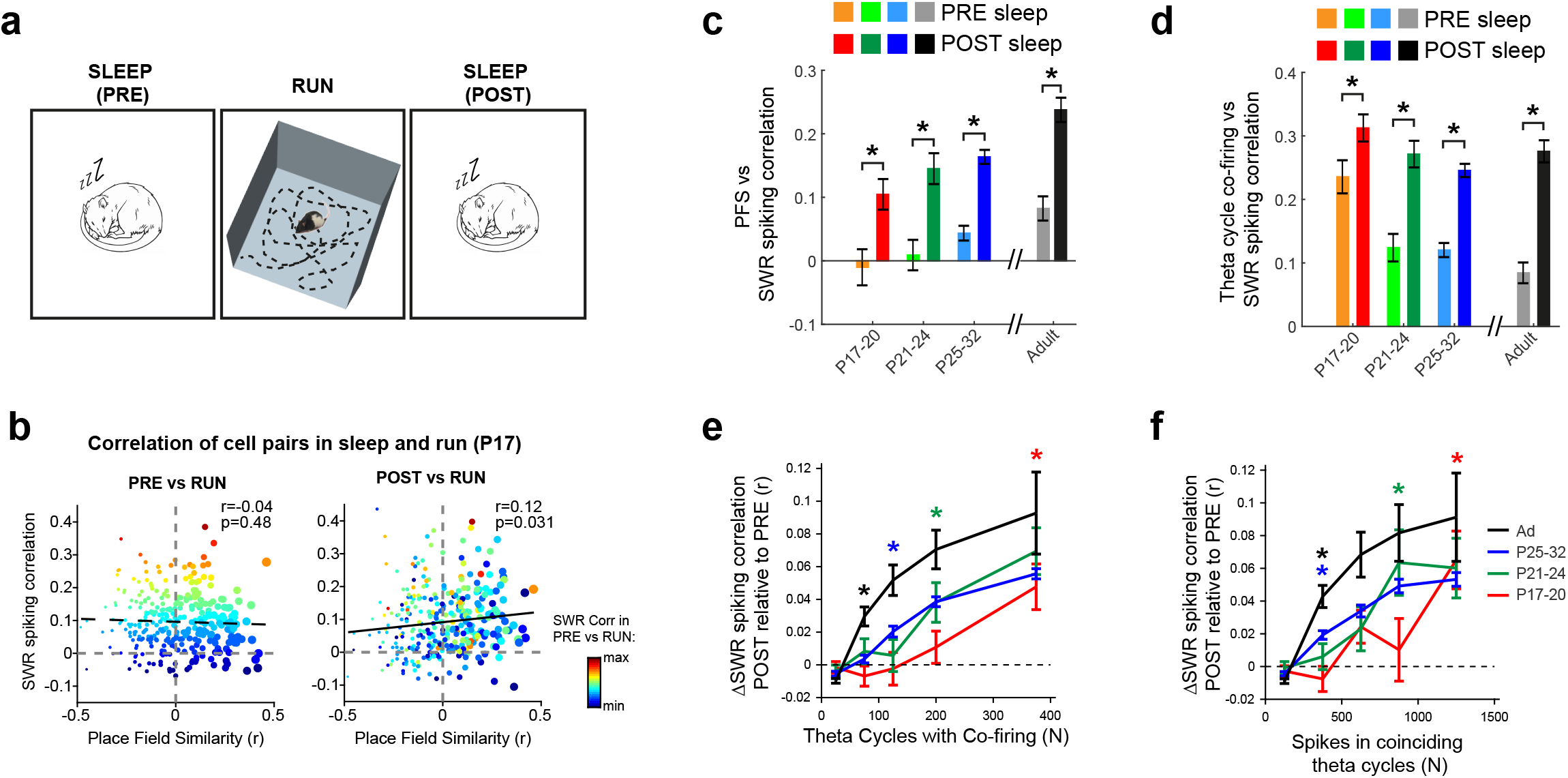
Reactivation of cell pair co-firing patterns during POST-experience sleep is already present at P17, but the amount of RUN co-firing required to induce plasticity is greater in young rats. (**a**) Schematic of experimental paradigm. Rats explored a square open field during RUN, and rested in a separate holding box in the same room before (PRE-sleep) and after (POSTsleep) open field exploration. (**b**) Example of cell pair co-activity correlation between RUN and temporally adjacent sleep sessions in one simultaneously recorded ensemble. Data were recorded at P17 and contain all cell pairs of the ensemble. X-axes show place field similarity in RUN (PFS; Pearson’s-r correlation of rate map bin values), y-axes correlation of co-firing during SWR events (correlation of cell-pair activity across all SWRs in sleep) in PRE (left panel), or POST (right panel), sleep sessions. Points are coloured according to magnitude of SWR spiking correlation in PRE and scaled in size according to their PFS in RUN. Regression statistics are in top right corner. Cell pairs with high PFS show an increase in SWR co-firing, during POST-sleep. (**c**) Bar chart showing Pearson’s-r values (±SE of correlation) of place field similarity (RUN) and SWR spiking correlation for PRE (pale colours) and POST (bold colours) sleep sessions, for all recorded cell pairs across development. Asterisks indicate differences at p≤0.001. (**d**) Bar chart showing Pearson’s-r values (±SE of correlation) of theta cycle co-firing (RUN) and SWR spiking correlation for PRE (pale colours) and POST (bold colours) sleep sessions across development. Asterisks indicate differences at p≤0.001. (e-f) Cell pair plasticity (change in cell-pair SWR spiking correlation from PRE to POST sleep) as a function of cell pair co-firing in RUN. (**e**) Cell pair plasticity as a function of the number of theta cycles in which both cells fire during RUN. (**f**) Cell pair plasticity as a function of the number spikes fired in theta cycles in which both cells fire. For (e-f), coloured asterisks mark the smallest x-axis bin in which cell pair plasticity is significantly different from zero (T-test of mean against 0; p<0.05), at each age.

In order to define the mechanism underlying reactivation during development, we tested how cell pair plasticity (defined as the change in SWR co-firing correlation from PRE– to POSTsleep sessions, for each neuron pair) depends on the degree of co-firing within theta cycles, during the exploratory phase (RUN) [22]. We found that, although increased RUN co-firing results in increased cell-pair plasticity at all ages, cell pairs in younger rats required significantly more RUN co-firing for plasticity to occur (Figure 1e, f). These data indicate that the co-activity threshold for cell-pair plasticity in the young hippocampus is higher than that in the adult, a result that is consistent with reports of heightened thresholds for the induction of long-term potentiation before P21 [23–25] (possibly compounded by the need to overcome stronger internal network dynamics in the youngest rats, as demonstrated by high correlations between PRE-sleep and RUN, see Figure 1d).

Overall our results demonstrate the existence of reactivation (increased sleep co-firing between pairs of neurons which were co-active during the preceding exploratory phase) in young rats. In addition to reactivation, in adult rats, the reinstatement of hippocampal network activity during off-line periods includes the ‘replay’ of temporally ordered sequences of neuronal firing which faithfully recapitulate the sequential firing observed during the exploratory phase. To investigate the emergence of replay during development, we recorded hippocampal neuronal activity in a sub-set of the rats which underwent reactivation testing, as they ran on a square corridor track in a familiar environment (Figure 2a; 1007 CS cells, 25 unique sessions from 16 rats, mean number of CS cells per session = 40.3, range 27-58). CS cells in young rats displayed spatially localised firing during locomotion (Figure 2b, as in[18,19,21]) with uniform distributions of CS cell firing across the environment (Figure 2c-d).

**Figure 2.**
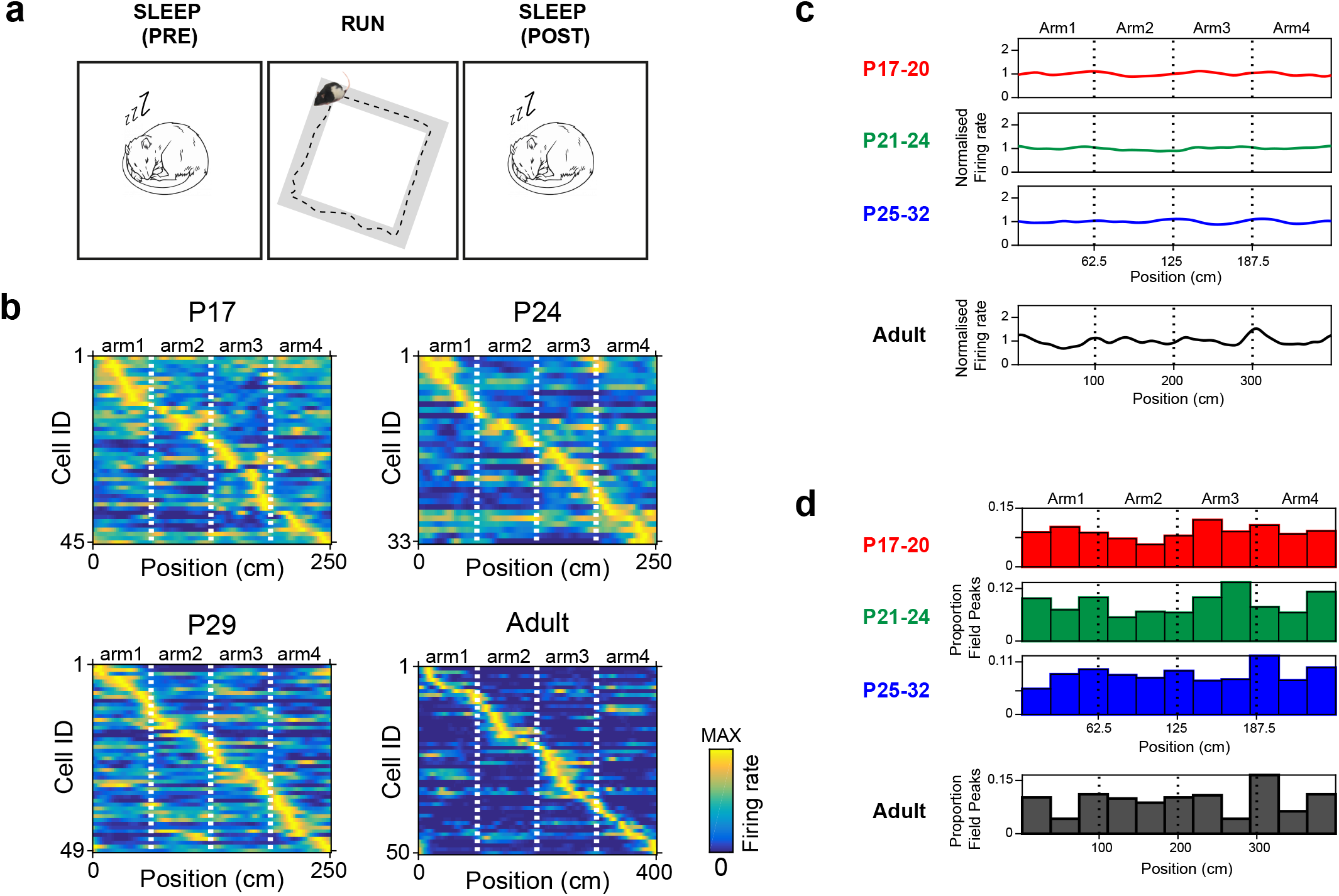
Complex spike cell firing on square track environment in developing rats. (**a**) Schematic of experimental paradigm. Rats explored a square track during RUN, and rested in a separate holding box in the same room before (PRE-sleep) and after (POST-sleep) track exploration. (**b**) Example place field maps for RUN sessions on square track at different ages. For each age, each row represents the spatial firing of one cell along the length of the square track, filtered for one running direction. False colours show firing rate, scaled to the peak firing rate for each cell. Cells are ordered according to position of peak spatial firing on the track. (**c-d**) CS cell spatial firing evenly covers the extent of the square track. (**c**) Mean spatial distributions of normalised firing rate of all CS cells recorded within each age group. (**d**) Histograms showing the proportion of CS cell peak spatial firing locations at different positions on the square track, within each age group.

In order to detect replay during sleep, we first defined ensemble spiking events as bursts of multi-unit activity (MUA) which coincided with SWRs (see Methods, Figure S1 for characterisation of sleep, SWR and MUA events). We then used Bayesian decoding followed by line fitting of time-by-position probability posteriors[26] to identify the presence of spiking representing linear trajectories through space. Importantly, for some of the linear fits the slope approximated zero, constituting events which contained place cell firing representing a single location on the track (from now on referred to as ‘stationary events’).

During the POST-sleep session, more events than those expected by chance exhibited linear trajectories, at all ages (Figure 3a, b; significance assessed by comparing each event to 500 cell identity-shuffled events; see Methods). Significant events in younger rats (<P21) were predominantly stationary, covering little or no distance on the track (Figure 3a, top four examples). The mean distance covered by linear trajectory events (Figure 1c, e) and their mean speed (Figure 1d, f) both gradually and linearly increased during development. Changes in event duration with age did not explain the increase in trajectory distance (Figure S2a-c). Developmental changes in linear trajectory distance are not caused by increases in the animals’ running speed, the length or duration of runs in a single direction or place cell spatial tuning, nor the extent to which decoded positions can be approximated by a linear trajectory (Figure S2d-i). Significant events appeared evenly distributed along the track, at all ages (Figure S3d). A gradual, linear, increase in event trajectory distance and speed was also observed when using an alternative method to determine event significance (rate map shift[26]; Figure S3a-c). Overall, our results demonstrate the gradual emergence of ordered place cell sequences during sleep SWRs (replay) between P17 and P32 in the rat, with the young hippocampus only capable of recalling single locations, and only gradually acquiring the ability to ‘stitch’ together separate locations into ordered trajectories spanning across space.

**Figure 3:**
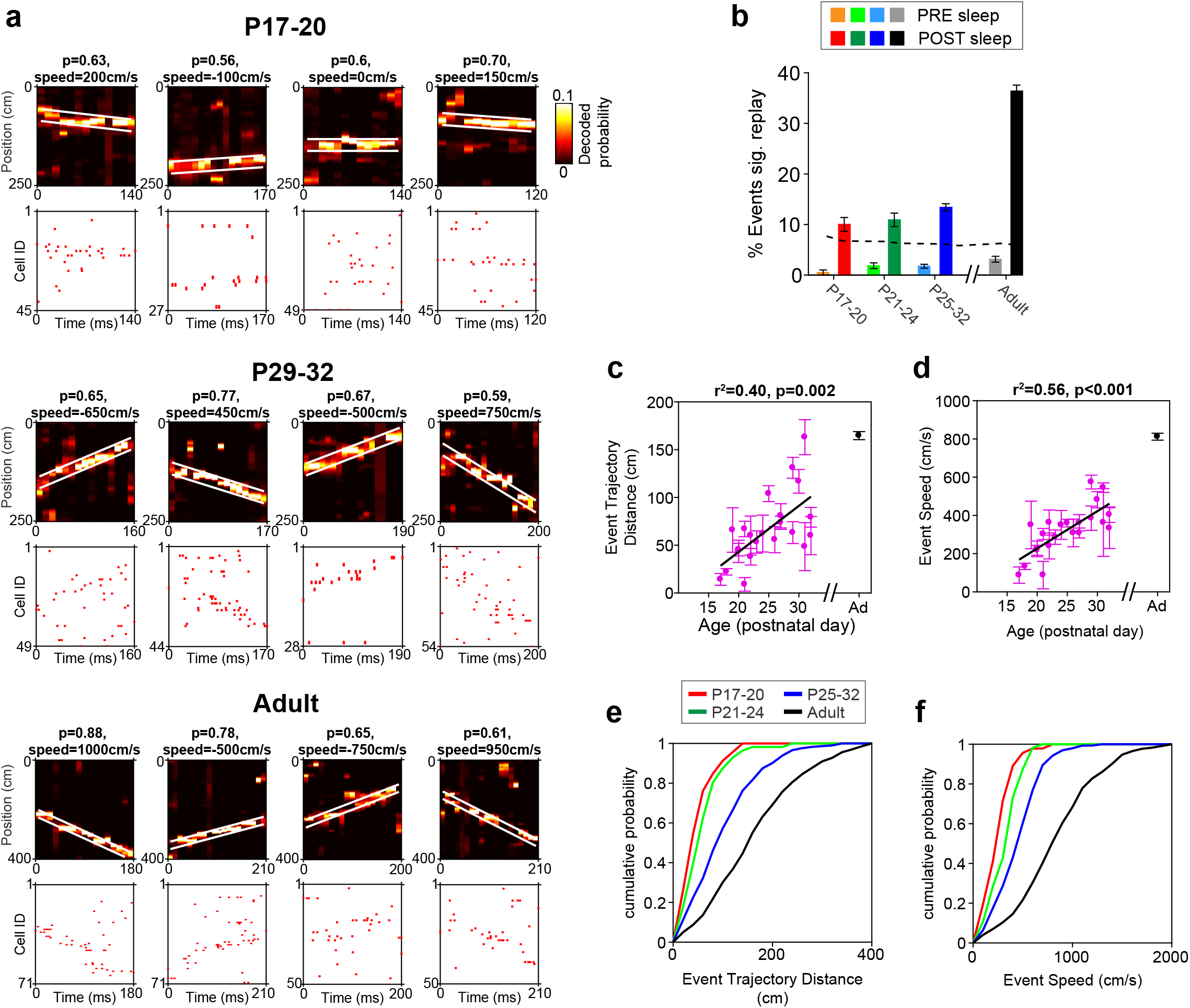
Gradual emergence of replay between P17 and P32. (**a**) Significant linear trajectory events in POST-sleep at different ages (four examples per age). For each event, top panel shows time-by-position probability posterior derived from Bayesian decoding of position, based on event spiking. False colours show decoded probability, white lines indicate the band of the best linear fit. Summed probability within fit lines (**p**) and speed of event (speed) are indicated above the posteriors. Bottom panel shows spike raster of complex cell activity during replay events. Cells are ordered by the position of their spatial peak firing on the track. Linear trajectories are predominantly stationary at younger ages, with replay emerging gradually in older animals. (**b**) Percentages (±95 confidence interval) of events with a significant linear trajectory, during PRE– and POST-sleep sessions across development. Dotted line represents 95% confidence threshold. In all age groups significantly more events than expected by chance showed a significant linear trajectory in POST sessions (binomial test, p<0.001 for all groups). (**c-d**) Mean characteristics of significant linear trajectory events in each POST-sleep sessions. For all plots, each data point represents mean (±SEM) of all significant linear events in one experimental session (one rat/day). Adult data represent overall mean across all sessions. For each measure r^2^ and p-values of linear regression over age is indicated above plots (adult data is always excluded from regression analysis). (**e**) Distance covered and (**f**) speed of decoded trajectories. (**e-f**) Cumulative distributions of the distance covered (**e**) and the speed (**f**) of all significant linear trajectory events, in the age groups P17-20, P21-24, P25-32 and in adult animals.

We investigated when theta sequences emerge during development by recording place cell activity in young rats during exploration of the square track (RUN trial). We then used Bayesian decoding to test whether the location encoded by the active CS population, relative to the actual location of the rat, varied within each theta cycle [27]. In adults and older pups, the encoded location shifts from behind the animal’s current position early in the theta cycle, to ahead of current position later in the theta cycle (Figure 4a). By the end of the 4^th^ week of life, therefore, CS cell firing during the theta cycle is organised into sequences defined by the relative locations of spatial firing fields on the track. However, qualitative examination of encoded positions relative to theta in younger pups reveals that theta sequences emerge slowly between P17 and P32 (Fig 4a; see Figure S4 for complete dataset). In order to quantify theta sequence occurrence we computed a ‘theta sequence score’, which captured systematic changes in decoded position relative to theta (see Methods). The theta sequence score increased gradually over the age range P17-32 (Figure 4b; correlation between theta sequence score and age: r^2^=0.54, p=0.001), confirming the qualitative impression that theta sequencing of place cell activity emerges gradually.

As new evidence suggests a link between theta sequence disruption during exploration and impaired replay sequences in subsequent sleep [28] in adult rats, we tested whether the theta sequence score in RUN was correlated with replay sequence incidence in POST-sleep during development. Strikingly, we observed that theta sequence score was strongly correlated with both replay trajectory speed and distance across all developmental ages (Figure 4c, d). Critically, these correlations hold even when age is controlled for (partial correlation of replay distance or speed with theta sequence score, controlling for age: distance; r^2^=0.36, p=0.004; speed; r^2^=0.19, p=0.048) and when overall mean firing rate and spatial information of complex spike cells are added as controlling variables (partial correlation, controlling for age, mean rate and spatial information: distance; r^2^=0.32, p=0.011; speed; r^2^=0.22, p=0.041). The partial correlations (controlling for age) are also significant when the RUN data is sub-sampled to match median running speeds across ages (distance: r^2^=0.36, p=0.004; speed: r^2^=0.19, p=0.048). Overall, these results demonstrate the coordinated emergence of theta and replay sequences during hippocampal post-natal development in the rat. Ordered sequences of hippocampal firing which occur during different brain states (theta sequences and replay, theta vs SWR) appear to be functionally linked early in development as much as in adulthood [28].

**Figure 4.**
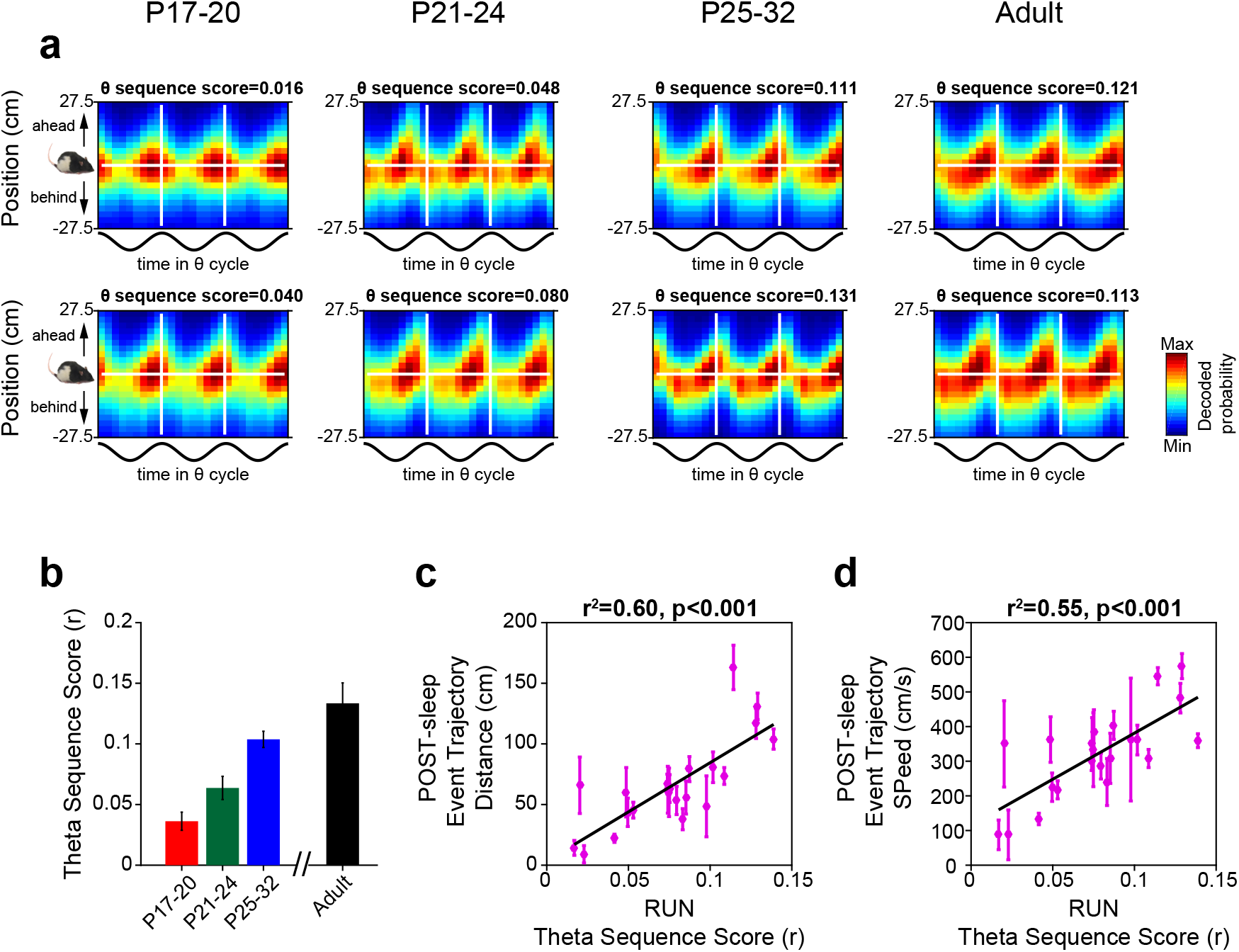
The gradual maturation of theta sequences between P17 and P32 is correlated with the emergence of replay. (**a**) Examples of theta sequence emergence across development. Each plot shows a probability posterior derived from a single RUN session, where the x-axis shows the proportion of time elapsed during the theta cycle and the y-axis shows position on the track relative to the current location of the rat. The horizontal white line shows current rat location, the vertical white lines demarcate one theta cycle. Hot colours show high decode probabilities. Numbers above the plots show theta sequence score, defined as the circular-linear weighted correlation of the probability posterior. Theta sequences are indicated by a shift in the decoded position from behind to ahead of the rat within the theta cycle: this emerges gradually between P17 and P32. (**b**) Mean (±SEM) theta sequences scores in each age group. (**c-d**) Theta sequence scores are correlated with the distance covered (**c**) and speed (**d**) of replay trajectories during development. Each data point represents mean (±SEM) of all significant linear events in one experimental session, for all developing rats. For each measure r^2^ and p-values of linear regression over age is indicated above plots. Correlations reported in (c, d) remain significant even after controlling for age, see main text.

## DISCUSSION

Here we have shown that reactivation of hippocampal activity in the open field – changes in cell-pair co-firing in sleep following exploration – occurs at the earliest ages tested (P17 onwards; Figure 1b-d) and that the activity of hippocampal neurons representing single locations (stationary linear trajectories) is reinstated during sleep immediately following exploration. In contrast, temporally ordered sequences of hippocampal firing describing extended trajectories through space (replay) emerge only gradually during development (Figure 2). Taken together these results indicate that functional assemblies encoding discrete locations are generated in the young hippocampus and selectively reactivated during off-line periods, thus excluding a generalised lack of Hebbian plasticity as the underlying cause of the developmental deficit in sequential replay in the young rodent. Interestingly, we found that in younger animals, the amount of cell pair co-activity required to express reactivation during sleep is higher in young animals (Fig 1e, f) indicating that a raised plasticity threshold in the young hippocampus may underlie the delayed emergence of sequential firing and spatial memory in young rodents [14,24]. According to this account, this increased threshold would be responsible for the predominance of stationary linear trajectory events in younger pups: only cells with very similar place fields are subject to plasticity in the developing hippocampus, reducing, for example, asymmetric plasticity [29] at the edges of widely spaced fields.

Hippocampal replay has been linked to cognitive functions such as memory consolidation, route planning and reward processing [30–32], in rodents as well as in humans [33]. Indeed, a computationally appealing feature of replay is that it could be used to chain together disparate spatial locations (or other, non-spatial events), without the requirement for the neurons representing those states to be co-active during exploration [34,35]. Here we have shown that sequenced firing during sleep, representing long trajectories through explored space, emerges gradually across the period P17 to P32 in the rat. This makes replay the latest emerging signature of hippocampal activity studied so far [18,19,21,36–38]. Interestingly, the timeline of replay development (slowly improving over the 4^th^ week of life) is a good match for previous reports of the emergence of spatial learning and memory in the rat [39].

We have also demonstrated the parallel, late and gradual emergence of theta sequences during development. Theta sequences are ordered sequences of hippocampal place cell firing which are nested within each theta cycle, and reproduce, at compressed timescales the spatial organisation of place fields in the environment. Critically, the emergence of theta sequences during exploration and replay during sleep appear to be strongly coordinated during development. These two phenomena are very likely functionally linked as they appear to be correlated not only in adulthood [28], but, as demonstrated here, from their first inception during development. Significantly though, replay during sleep (POST-sleep) does not simply recapitulate theta sequences during the preceding exploratory phase (RUN), as the length of replay trajectories exceeds that of theta sequences (e.g. approximately 30cm for theta sequences and 1m for replay, in older pups). In adult rodents, theta sequences require plasticity to emerge [27], possibly reliant on functional inputs from CA3 circuits [40]; similarly, there is evidence indicating that replay requires NMDA-R dependent plasticity [41]. The heightened threshold for cell pair plasticity we observed in young rodents may therefore be the underlying cause for the delayed and coordinated emergence of both theta and replay sequences during development.

Altered replay of place cell sequences has previously been reported in aging and rodent models of cognitive impairment [42,43], strongly suggesting the existence of a functional link between replay and the ability of the hippocampus to successfully encode and store memory traces. Exploiting the natural emergence of hippocampal function during development, our study shows, for the first time, that, during development, the appearance of sequential firing representing extended trajectories (spatial or otherwise), may be a pre-requisite for the emergence of its mnemonic and navigational functions.

## Methods

### Subjects

24 male Lister Hooded rat pups, aged P13-P24 and weighing 26-69 g at time of surgery were used as subjects. Litters were bred in-house and remained with their dams until weaning (P21). Rats were maintained on a 12:12 hour light:dark schedule (with lights off at 10:00). At P4, litters were culled to 8 pups per mother in order to minimise inter-litter variability. Pregnant females were checked at 17:00 daily and if a litter was present, that day was labelled P0. After surgery (see below), each pup was separated from the mother for between 30 minutes and 2 hours each day, to allow for electrophysiological recordings. 2 male Lister Hooded adult rats, aged 3-6 months at the time of recording, were included in the study to provide a comparison for the pup data.

### Surgery and electrode implantation

Rats were anaesthetised using 1 -2% isoflurane, and 0.15mg/Kg bodyweight buprenorphine. Rats were chronically implanted with microdrives loaded with 8-16 tetrodes (HM-L coated 90% platinum-10% iridium 17μm diameter wire). Microelectrodes were aimed at the hippocampal CA1 region, using the co-ordinates 2.9 mm posterior and 1.8 mm lateral to Bregma. After surgery, rats were placed in a heated chamber until they had fully recovered from the anaesthetic (10 – 30 minutes), and were then returned to the mother and littermates.

### Single-unit recording

Rats were allowed a 1-day postoperative recovery, after which microelectrodes were advanced ventrally by 62-250 μm/day until they reached the hippocampal CA1 pyramidal cell layer, identified physiologically by the presence of complex spike cells and 140-200Hz ‘ripple’ fast oscillations. When CA1 complex spike (CS) cells were detected, recording sessions began. Single units recorded in the CA1 pyramidal cell layer were defined as CS cells (putative pyramidal cells) using the following criteria: a) spike width (from peak to following trough) ≥ 300ms, b) first moment of the temporal autocorrelogram of the cell (within a 50ms window) ≤25ms, c) mean firing rate ≤ 5Hz. Single unit data was acquired using an Axona (Herts, UK) DACQ system. LEDs, were used to track the position and directional heading of the animal. Isolation of single units from multi-unit data was performed manually on the basis of peak-to-trough amplitude, using the software package ‘TINT’ (Axona, Herts, UK). Isolated single units were only used for further analysis if they fired 75 spikes or more within a RUN trial.

### Behavioural testing

During RUN trials (awake exploration), single-unit activity was recorded while rats searched for drops of soya-based infant formula milk randomly scattered on the floor of two different environments. Open field RUN trials used a square-walled (62.5cm sides, 50cm high) light-grey wooden box, placed on a black, square platform. Square track RUN trials used the same environment, with the addition of a matt gray wooden insert, 25 cm height, which constrained rats to run on a 8.5 cm track in the vicinity of the walls. For adults, a larger square track (of similar construction; height of inset = 45cm) was used, with 1m arms and a 10.5 cm track width. Reward was scattered pseudo-randomly, to encourage rats to run in both clockwise (CW) and counterclockwise (CCW) directions around the track. Pilot experiments showed that this track configuration resulted in a more consistent running than in a linear runway track, in which young rats showed a tendency to sit at the track ends. Trials were 15 minutes long. Distal visual cues were available in the form of the fixed apparatus of the laboratory. During PRE and POST sleep trials, rats were kept in a separate holding box (30×30cm) in the same room. No intra-maze cues from the RUN environment were visible from the sleep box.

### Construction of Firing Rate Maps: open field

All spike and positional data were filtered so as to remove periods of immobility (defined as speed < 2.5cm/s). For the open field, positional data were then sorted into 2.5 × 2.5 cm spatial bins. Following this, total positional dwell time and spike count for the whole trial was calculated for each spatial bin. The binned position dwell time and spike count maps for each cell were then smoothed using adaptive smoothing[1]. In brief, to calculate the firing rate for a given bin, a circle centred on the bin is gradually expanded in radius *r* until

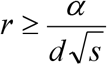

where α=200 and *d* and *s* are the positional dwell time (in seconds) and the number of spikes lying within the circle, respectively. The firing rate assigned to the bin is then set equal to *s/d*.

### Construction of Firing Rate Maps: square track

For square track trials, position data was linearised, by radially binning positions with a set of bin edges corresponding to points evenly spaced (0.25cm apart) along each arm of the square track. Following linearization, the position data was sorted into CW and CCW runs, defined as 5s epochs of constant running (excluding immobility) in either direction. Two sets of rate maps, CW and CCW, were then constructed, by re-binning the linearized positional and spike data into larger, 2.5cm long bins, and taking the overall summed dwell and spike count in these bins. The spike and dwell maps were then smoothed with a Gaussian kernel (s.d. 5cm) before the overall rate maps were made, by dividing summed spikes by summed dwell, in each bin.

### Detection of slow-wave sleep, sharp-wave ripples and multi-unit activity bursts

The brain states slow-wave sleep (SWS), rapid-eye movement sleep (REM) and awake movement were defined following[2]. A multitaper power spectral density estimate of the hippocampal local field potential (LFP) was derived for 1.6s windows, overlapping by 0.8s (matlab function ‘pmtm’). From this, power in the delta and theta bands were calculated in each window. As theta frequency changes during development[3], theta and delta peak frequencies were calculated for each session, defined as the peak frequency of the fast Fourier transform of the LFP, in the bands 5-11Hz (theta) and 1.5-4Hz (delta). Mean running speed for each 1.6s bin was also estimated. Slow-wave sleep was then defined as epochs with speed <2cm/s, and theta/delta power ratio <2, REM as epochs with theta/delta power ratio >2 and speed <1cm/s, and waking movement as theta/delta power ratio >2 and speed >2.5cm/s. Sharp-wave ripples were detected by first filtering the LFP in the band 100-250Hz. The instantaneous power of the filtered LFP was then estimated by calculating the root mean square over 7ms intervals (matlab function ‘envelope’ with option ‘rms’). From all LFPs across tetrodes in the CA1 layer, the power estimate with the highest standard deviation was then used to define ripple events, as 100ms windows around the peak power, whenever the power was greater than the 99^th^ percentile of all powers in the trial (approximately equal to 4 standard deviations above the mean). Multi-unit activity (MUA) bursts were defined by binning all complex spikes into 1 ms bins and smoothing the resulting binned spike train with a Gaussian kernel (s.d. 5ms). MUA events were then defined as crossing of a threshold defined as 3 standard deviations above the mean of the smoothed spike train, with a duration from 100-500ms. Only MUA bursts which temporally overlapped (even in part) with SWR events were included in the replay analysis.

### Reactivation analyses

SWR cell pair correlations were calculated by, for each cell, binning all spikes occurring in SWR windows, during SWS epochs. The SWS cell pair correlation was then defined as the correlation of the two binned spike count vectors[2]. Place field similarity was defined as the correlation (Pearson’s-r) between the rate values of spatially corresponding bins, in the two rate maps. To define individual theta cycles (for RUN analyses), theta phase for each LFP sample was first defined using the Hilbert transform. The preferred ensemble phase was defined, for each session, as the theta phase with the highest ensemble firing, and theta cycle bin edges were defined as phase crossings 180° out of phase with the preferred ensemble phase. Following theta cycle definition, spikes for each cell were binned into theta cycle temporal bins, and theta cycle correlation was then defined as the correlation of the two binned spike count vectors. Only theta cycles from ‘waking movement’ brain state epochs were used (see above). Counts of N theta cycles with co-firing and N spikes in coinciding theta cycles were defined using the same set of theta cycle temporal bins. All 24 developing rats contributed to the reactivation analysis, in 44 unique experimental sessions, yielding a total of 19,334 cell pairs.

### Replay analysis

Spiking in SWR/MUA joint events was split into overlapping 20ms temporal bins (overlap 10ms), spanning the duration of the MUA burst. For each temporal bin, the probability of dwelling in each spatial bin of the linear rate map was then estimated using a Bayesian probability framework, as described in[4]. To estimate the goodness-of-fit of the decoded posterior to a linear trajectory, the summed probability under a linear band, 25 cm wide, was maximized by searching the set of all lines starting every 2.5cm along track, and with a range of line slopes corresponding to 0 – 2500cm/s, in steps of 50cm/s. As the square track has a circular topology, linear bands were wrapped around the posterior when they reached its first or last spatial bin. The fitting procedure was run independently for both CW and CCW maps, and the best overall fit was taken as the final trajectory. The likelihood of this fit was then determined by comparing the actual best fit to a population of 500 fits of posteriors based on shuffled data, for each event. Only events whose best summed probability exceeded the 95^th^ percentile of the shuffled events summed probabilities were defined as exhibiting a significant linear trajectory. Shuffled data was produced by either randomizing cell identities (of MUA spiking with respect to rate maps; ‘cell identity shuffle’) or subjecting each rate map to a different random linear offset (‘spatial shuffle’), before performing decoding. Shuffling for each of the CW and CCW rate map sets was performed independently, and the best fit across directions, for each event, contributed to the final shuffled population. Shuffling was therefore not yoked to the rate map direction of the actual posterior best fit[5]. The speed of replay trajectory was defined as the slope of the best fit, and the distance covered defined as the speed multiplied by the MUA event duration. Replay analysis was only applied to CS cell ensembles in which >25 CS cells fired >75 spikes during RUN (after immobility filtering was applied), and where the ‘online’ Bayesian decoding of RUN positions from RUN spiking[6] (in 300ms time windows) predicted the current position of the rat with an overall median error <10cm. 25 unique ensembles from 16 developing rats passed these criteria.

### Theta Sequence Analyses

Following[7], Bayesian decoding was applied to RUN data, using 20ms decode windows, sliding in 5ms steps. Only data epochs of >5s constant running in one direction, at speeds >=2.5cm/s was used. The resulting probability posterior for each temporal decode window was then shifted such that the actual position of the rat during the window was at 0 cm. Posteriors for runs in one direction only were then reversed, such that the reference frame for the whole posterior represented position relative to current position and direction of travel. Theta cycles were demarcated as described above (‘*Reactivation analyses*’) and for each decode window, the corresponding time elapsed within the current theta cycle was determined by linear interpolation between the times of the two peaks. Decode windows were then binned into 10 bins, each representing an equal proportion of the time elapsed through the theta cycle, and the overall mean posterior probability was calculated for each theta cycle time bin. To exclude the effect of running speed on the development of theta sequences, the analysis was repeated with RUN data that was sub-sampled to match median run speeds across sessions and ages. To do this, decode windows were sorted in order of their corresponding running speeds, and either fast or slow decode windows were then discarded, in order of speed, until the median decode window for the trial was equal to 10.6cm/s (overall median running speed for dataset, after exclusion of data <2.5cm/s).

### Circular-linear weighted correlation of theta sequences

Previous approaches to quantifying theta sequences [7,8] were not well-suited to developmental data: the preferred theta phase of CS cell firing changed during development (see Fig 4a) meaning that it was not possible to define a consistent set of phase by position ‘quadrants’ on the theta sequence, which would be needed for a ‘quadrant score’ [7]. Furthermore, maturing theta sequences did not always approximate a linear band, meaning that a linear band fitting approach [8] could not be used. Instead, the tendency of decode probabilities to systematically change throughout the theta cycle was assessed by correlating position and theta cycle time bin, weighted by the posterior probability for each bin (analogous to the weighted correlation of a probability posterior in [5]). However, to take account of the circular nature of the theta cycle, the weighted correlation was circular-linear. A standard circular-linear correlation coefficient, *r_cl_*, can be computed as:

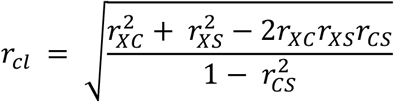

Where *r_XC_* is the (linear) correlation coefficient between the linear variable and the cosine of the angular variable, *r_XS_* is the (linear) correlation coefficient between linear variable and the sine of the angular variable and *r_CS_* is the (linear) correlation coefficient between the sine and cosine of the angular variable[9]. To calculate the weighted circular-linear correlation of the theta sequence probability posterior, the coefficients *r_XC_, r_XS_* and *r_CS_* were calculated by correlating position bin distance with theta bin angle, weighted by the corresponding p-value of the posterior. These coefficients were then used to calculate, *r_cl_*, following the equation above.

---

**Figure S1.**
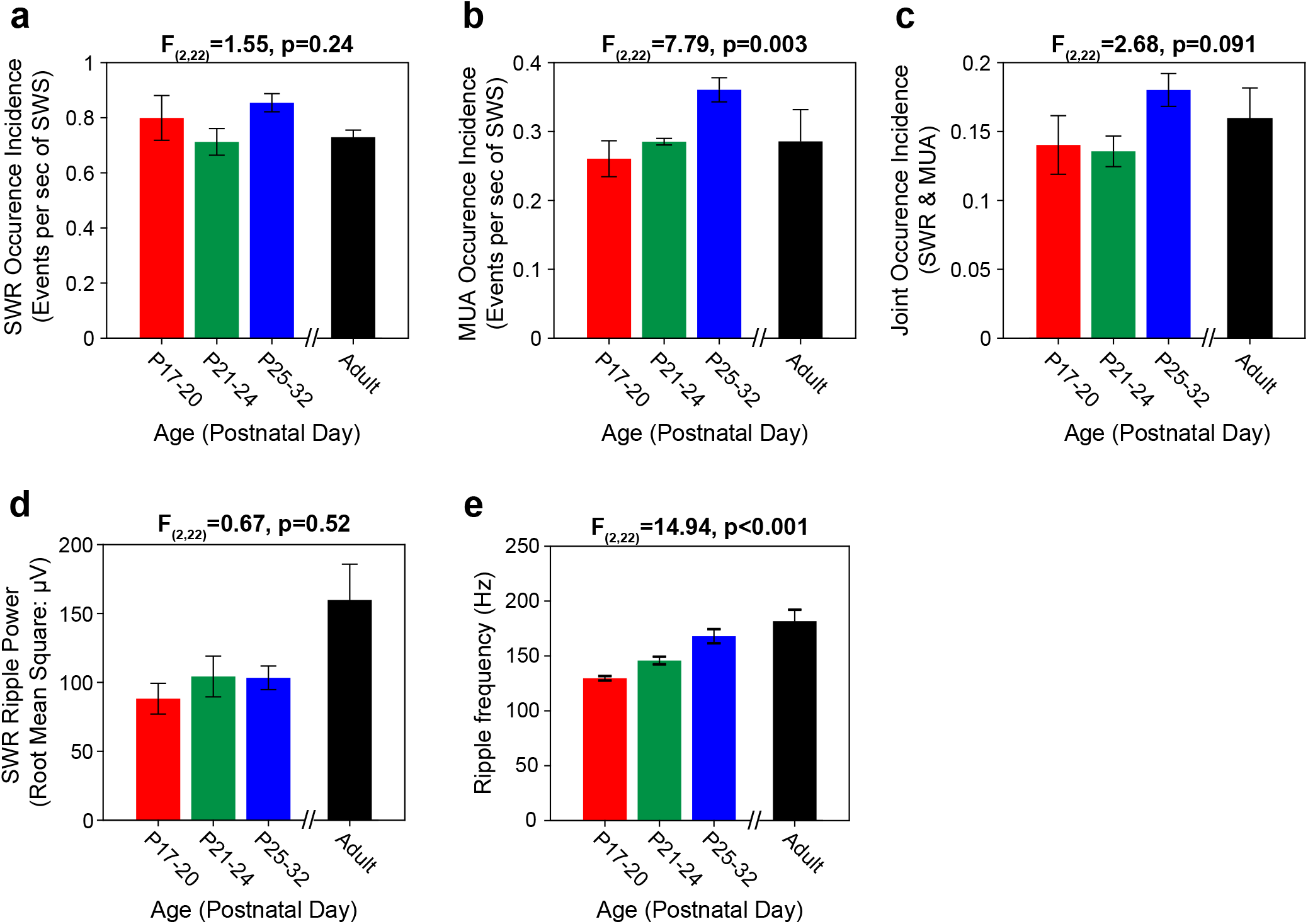
**related to figure 3**.Characterisation of SWRs and MUA bursts during development. (**a-c**) Bar charts showing the mean incidence (±SEM) of the occurrence of SWRs (a), MUAs (**b**) and overlapping SWR/MUA events (c), in each experimental session. All y-axes show events per second of SWS. 1-way ANOVA statistics comparing developing rat groups shown on top of graph. There is a significant increase in the occurrence of MUAs during development, but no significant change in the occurrence of SWRs, or overlapping SWR/MUA events. (**d**) Bar chart showing the average root-mean-square power (±SEM) of SWRs in each experimental session. There is no significant change in SWR power during development (1-way ANOVA statistics comparing developing rat groups shown on top of graph), though the power of adult SWRs is greater than that of those in developing rats. (**e**) Bar chart showing the mean ripple frequency (±SEM) of SWRs in each experimental session. There is a significant increase in ripple frequency during development: 1-way ANOVA statistics comparing developing rat groups shown on top of graph.

**Figure S2.**
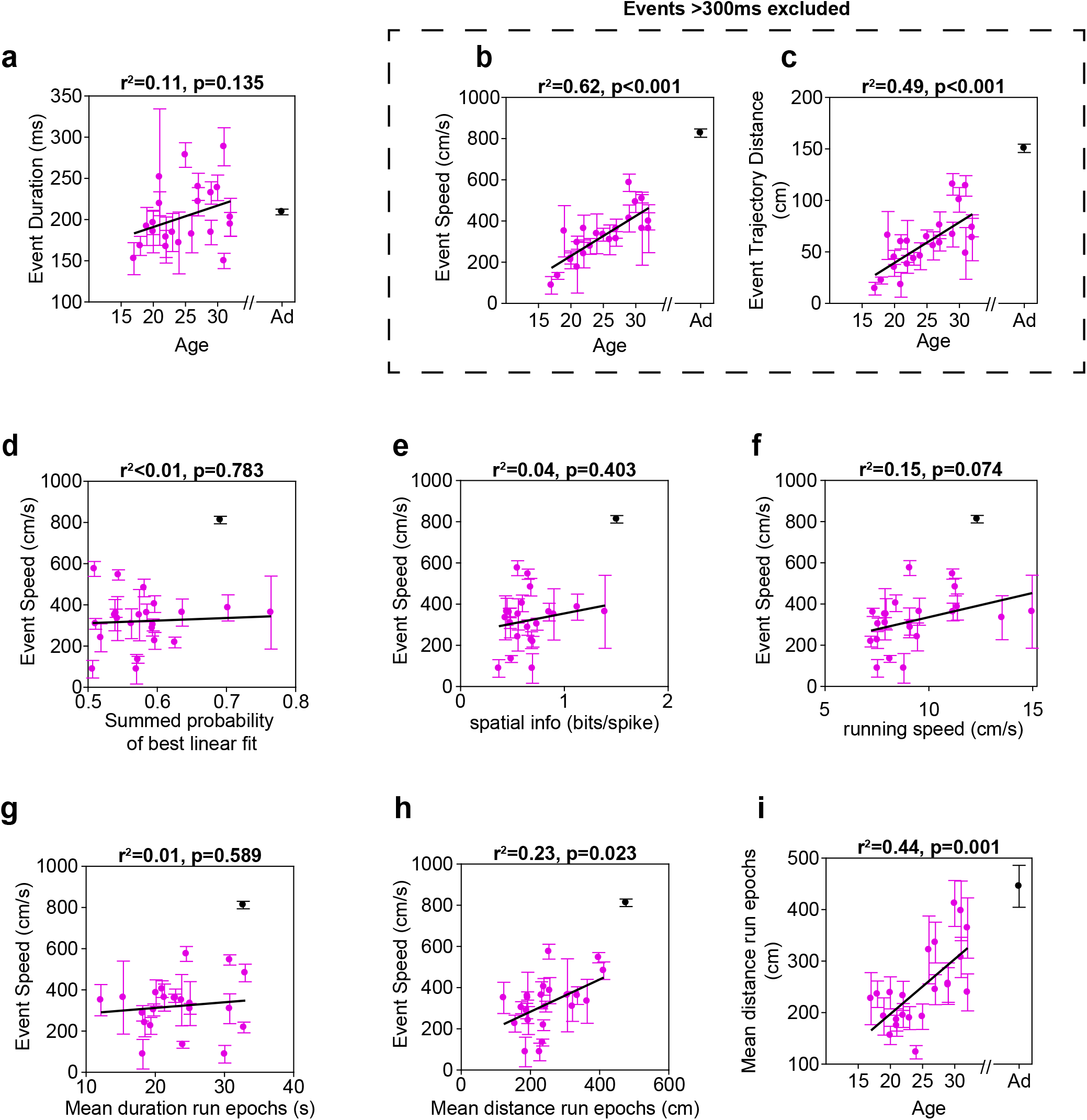
**related to figure 3**.Developmental increases in linear trajectory distance and speed of significant events are not due to changes in event duration, changes in place cell tuning, running speed or linear fitting. In all panels each pink data point indicates the mean of a characteristic of significant linear events of one developing rat at one age point (±SEM). Adult event mean is shown in black, and is excluded from correlations. Correlation statistics are shown above plots. (a) No significant correlation between average MUA event duration and age of animals. (b-c) Furthermore, increases in mean trajectory distance (b) and speed within each rat remain significantly correlated with age, even after events of >300ms are removed. (d) No significant correlation between the mean goodness of the linear fit (defined as the summed probability within the linear fit band) and the mean speed of linear trajectory events. (e) No significant correlation between mean linear trajectory event speed and the spatial tuning of complex spike cells (defined as the mean spatial information of the rate maps of the recorded ensemble, during exploration). (f) Trend for a significant correlation between mean linear trajectory event speed and the exploration speed of the rat (defined as the median running speed for the session, after exclusion of periods of immobility), as both these increase with age. The partial correlations of age versus event trajectory distance and speed, after controlling for running speed, remain significant (Distance r^2^=0.49, p<0.001, speed r^2^=0.52, p<0.001). (g) No significant correlation between the mean temporal duration of epochs of constant running direction in RUN sessions, and the mean speed of significant linear trajectories. (h-i) Significant correlation between mean distance traversed during epochs of constant running direction and the mean speed of significant linear trajectory events (h) as both scores increase with age (see (i) for scatterplot of mean distance traversed during epochs of constant running and age). However, the partial correlation for age versus distance and speed, after controlling for epoch distance, remain significant (distance r^2^=0.22, p=0.033, speed r^2^=0.42, p=0.001).

**Figure S3.**
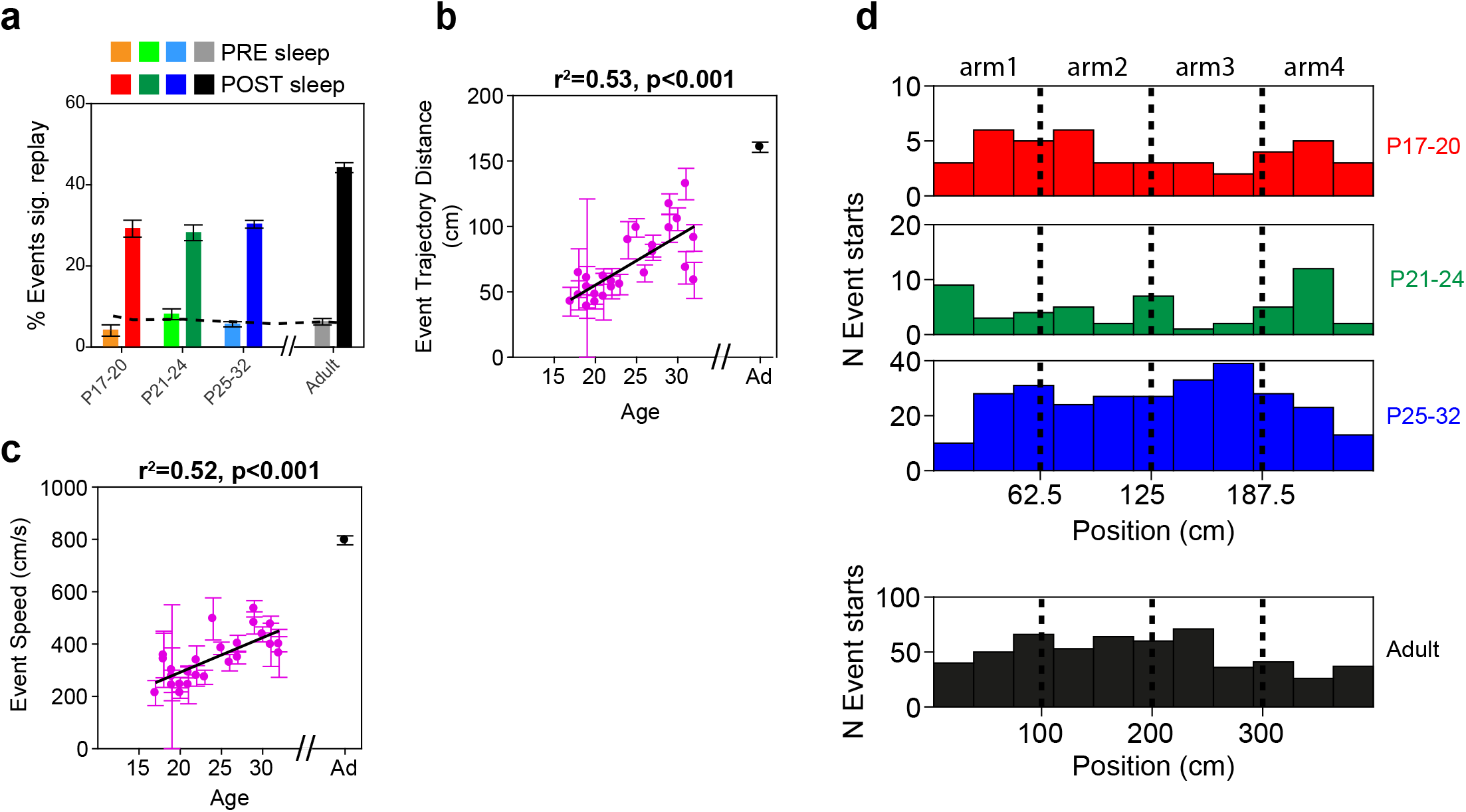
**related to figure 3**. Gradual emergence of replay between P17 and P32: results are unchanged when significance of event linear fit is determined by comparison to a population of shuffled events generated by randomly shifting spatial firing rate maps before decoding (‘map shuffle’). In all of (b-c) each pink data point indicates the mean of a characteristic of significant replay events of one developing rat at one age point (±SEM). Adult event mean is shown in black, and is excluded from correlations. Correlation statistics are shown above the plots. (a) Bar charts showing the percentage of spiking events determined to contain significant linear trajectories following the map shuffle, in both PRE and POST sleep, at all ages. Bars show the 95% confidence intervals of the percentages, the dashed line indicates the 95% confidence level. Note that more events in developing rats are classified as containing significant linear trajectories, compared to the cell identity shuffle. (b-c) Mean distance covered (b) and mean speed (c) of significant linear trajectory events when using the map shuffle to determine event significance. Bars show ±SEM of scores within rat. Both scores show significant correlations with age. (d) Significant linear trajectory events start at positions equally distributed throughout the square track. Histograms show counts of significant events with trajectories starting at corresponding points on the square track. At all ages, trajectories show no apparent bias to start at particular positions on the track.

**Figure S4.**
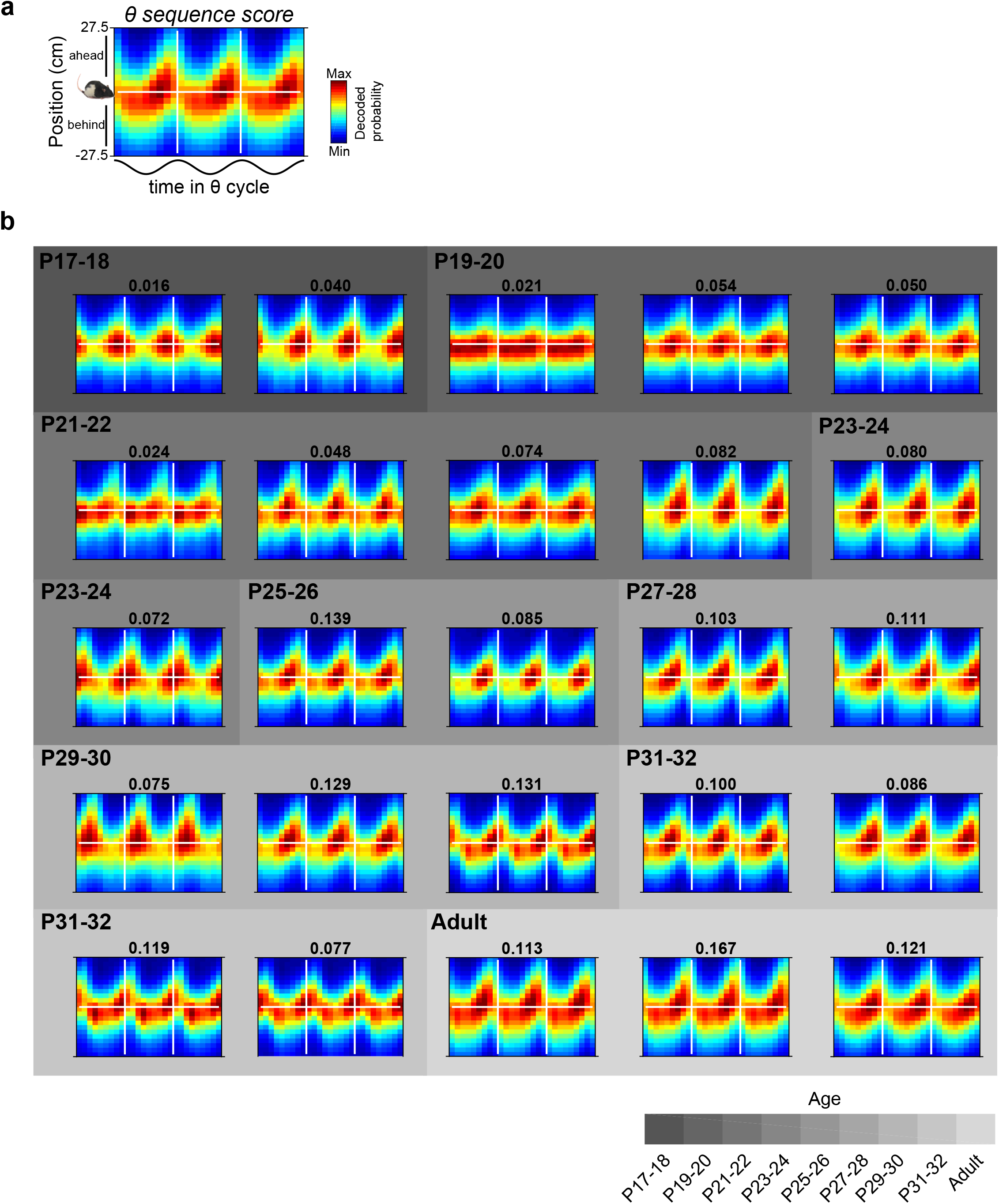
**related to figure 4**. Theta sequence probability posteriors for all datasets. (a) Schematic example exemplifying format of posteriors displayed in (b). The probability posterior is derived from a single RUN session, where the x-axis shows the proportion of time elapsed during the theta cycle and the y-axis shows position on the track relative to the current location of the rat. The horizontal white line shows current rat location, the vertical white lines demarcate one theta cycle. Hot colours show high decode probabilities. Numbers above the plots show theta sequence score, defined as the circular-linear weighted correlation of the probability posterior. Theta sequences are indicated by a shift in the decoded position from behind to ahead of the rat within the theta cycle: this emerges gradually between P17 and P32. (b) Theta sequence emergence across development. Probability posteriors of all datasets are shown, ordered by age. Age key for background grey shading of posteriors is shown below.

## References

1. O’Keefe, J., and Dostrovsky, J. (1971). The hippocampus as a spatial map. Preliminary evidence from unit activity in the freely-moving rat. Brain Res. 34, 171–175.

2. Wilson, M.A., and McNaughton, B.L. (1994). Reactivation of hippocampal ensemble memories during sleep. Science (80-.). 265, 676–679.

3. Skaggs, W.E., and McNaughton, B.L. (1996). Replay of neuronal firing sequences in rat hippocampus during sleep following spatial experience. Science (80-.). 271, 1870–1873.

4. Nadasdy, Z., Hirase, H., Czurko, A., Csicsvari, J., and Buzsaki, G. (1999). Replay and time compression of recurring spike sequences in the hippocampus. J.Neurosci. 19, 9497–9507.

5. Lee, A.K., and Wilson, M.A. (2002). Memory of sequential experience in the hippocampus during slow wave sleep. Neuron 36, 1183–1194.

6. Dupret, D., O’Neill, J., Pleydell-Bouverie, B., and Csicsvari, J. (2010). The reorganization and reactivation of hippocampal maps predict spatial memory performance. Nat. Neurosci. 13, 995–1002.

7. McNamara, C.G., Tejero-Cantero, Á., Trouche, S., Campo-Urriza, N., and Dupret, D. (2014). Dopaminergic neurons promote hippocampal reactivation and spatial memory persistence. Nat. Neurosci. 17, 1658–60.

8. van de Ven, G.M., Trouche, S., McNamara, C.G., Allen, K., and Dupret, D. (2016). Hippocampal Offline Reactivation Consolidates Recently Formed Cell Assembly Patterns during Sharp Wave-Ripples. Neuron 92, 968–974.

9. Girardeau, G., Benchenane, K., Wiener, S., Buzsáki, G., and Zugaro, M. (2009). Selective suppression of hippocampal ripples impairs spatial memory. Nat. Neurosci. 12, 1222–1223.

10. Ego-Stengel, V., and Wilson, M.A. (2009). Disruption of ripple-associated hippocampal activity during rest impairs spatial learning in the rat. Hippocampus 20, 1–10.

11. Dragoi, G., and Buzsáki, G. (2006). Temporal encoding of place sequences by hippocampal cell assemblies. Neuron 50, 145–57.

12. Foster, D.J., and Wilson, M.A. (2007). Hippocampal theta sequences. Hippocampus 17, 1093–1099.

13. Wikenheiser, A.M., and Redish, A.D. (2015). Hippocampal theta sequences reflect current goals. Nat. Neurosci. 18, 289–294.

14. Dumas, T.C. (2005). Late postnatal maturation of excitatory synaptic transmission permits adult-like expression of hippocampal-dependent behaviors. Hippocampus 15, 562–578.

15. Wills, T.J., Muessig, L., and Cacucci, F. (2014). The development of spatial behaviour and the hippocampal neural representation of space. Philos. Trans. R. Soc. Lond. B. Biol. Sci. 369, 20130409.

16. Alberini, C.M., and Travaglia, A. (2017). Infantile Amnesia: A Critical Period of Learning to Learn and Remember. J. Neurosci. 37, 5783–5795.

17. Mullally, S.L., and Maguire, E.A. (2014). Learning to remember: The early ontogeny of episodic memory. Dev. Cogn. Neurosci. 9, 12–29.

18. Wills, T.J., Cacucci, F., Burgess, N., and O’Keefe, J. (2010). Development of the hippocampal cognitive map in preweanling rats. Science (80-.). 328, 1573–1576.

19. Langston, R.F., Ainge, J.A., Couey, J.J., Canto, C.B., Bjerknes, T.L., Witter, M.P., Moser, E.I., and Moser, M.B. (2010). Development of the spatial representation system in the rat. Science (80-.). 328, 1576–1580.

20. Muessig, L., Hauser, J., Wills, T.J., and Cacucci, F. (2015). A Developmental Switch in Place Cell Accuracy Coincides with Grid Cell Maturation. Neuron 86, 1167–1173.

21. Scott, R.C., Richard, G.R., Holmes, G.L., and Lenck-Santini, P.P. (2010). Maturational dynamics of hippocampal place cells in immature rats. Hippocampus.

22. O’Neill, J., Senior, T.J., Allen, K., Huxter, J.R., and Csicsvari, J. (2008). Reactivation of experience-dependent cell assembly patterns in the hippocampus. Nat. Neurosci. 11, 209–15.

23. Dumas, T.C. (2012). Postnatal alterations in induction threshold and expression magnitude of long-term potentiation and long-term depression at hippocampal synapses. Hippocampus 22, 188–99.

24. Blair, M.G., Nguyen, N.N.-Q., Albani, S.H., L’Etoile, M.M., Andrawis, M.M., Owen, L.M., Oliveira, R.F., Johnson, M.W., Purvis, D.L., Sanders, E.M., et al. (2013). Developmental changes in structural and functional properties of hippocampal AMPARs parallels the emergence of deliberative spatial navigation in juvenile rats. J. Neurosci. 33, 12218–28.

25. Buchanan, K.A., and Mellor, J.R. (2007). The development of synaptic plasticity induction rules and the requirement for postsynaptic spikes in rat hippocampal CA1 pyramidal neurones. J. Physiol. 585, 429–445.

26. Davidson, T.J., Kloosterman, F., and Wilson, M.A. (2009). Hippocampal Replay of Extended Experience. Neuron 63, 497–507.

27. Feng, T., Silva, D., and Foster, D.J. (2015). Dissociation between the Experience-Dependent Development of Hippocampal Theta Sequences and Single-Trial Phase Precession. J. Neurosci. 35, 4890–4902.

28. Drieu, C., Todorova, R., and Zugaro, M. (2018). Nested sequences of hippocampal assemblies during behavior support subsequent sleep replay. Science (80-.). 362, 675–679.

29. Mehta, M.R. (2007). Cortico-hippocampal interaction during up-down states and memory consolidation. Nat. Neurosci. 10, 13–15.

30. Carr, M.F., Jadhav, S.P., and Frank, L.M. (2011). Hippocampal replay in the awake state: a potential substrate for memory consolidation and retrieval. Nat. Neurosci. 14, 147–153.

31. Foster, D.J. (2017). Replay Comes of Age. Annu. Rev. Neurosci. 40, 581–602.

32. Ólafsdóttir, H.F., Bush, D., and Barry, C. (2018). The Role of Hippocampal Replay in Memory and Planning. Curr. Biol. 28, R37–R50.

33. Kurth-Nelson, Z., Economides, M., Dolan, R.J., and Dayan, P. (2016). Fast Sequences of Non-spatial State Representations in Humans. Neuron 91, 194–204.

34. Foster, D.J., and Knierim, J.J. (2012). Sequence learning and the role of the hippocampus in rodent navigation. Curr. Opin. Neurobiol. 22, 294–300.

35. Mattar, M.G., and Daw, N.D. (2018). Prioritized memory access explains planning and hippocampal replay. Nat. Neurosci. 21, 1609–1617.

36. Wills, T.J., Barry, C., and Cacucci, F. (2012). The abrupt development of adult-like grid cell firing in the medial entorhinal cortex. Front Neural Circuits. 6, 21.

37. Bjerknes, T.L., Moser, E.I., and Moser, M.-B. (2014). Representation of geometric borders in the developing rat. Neuron 82, 71–8.

38. Muessig, L., Hauser, J., Wills, T.J., and Cacucci, F. (2016). Place Cell Networks in Pre-weanling Rats Show Associative Memory Properties from the Onset of Exploratory Behavior. Cereb. Cortex 26, 3627–3636.

39. Schenk, F. (1985). Development of place navigation in rats from weaning to puberty. Behav.Neural Biol. 43, 69–85.

40. Middleton, S.J., and McHugh, T.J. (2016). Silencing CA3 disrupts temporal coding in the CA1 ensemble. Nat. Neurosci. 19, 945–51.

41. Silva, D., Feng, T., and Foster, D.J. (2015). Trajectory events across hippocampal place cells require previous experience. Nat. Neurosci. 18, 1772–1779.

42. Gerrard, J.L., Burke, S.N., McNaughton, B.L., and Barnes, C.A. (2008). Sequence Reactivation in the Hippocampus Is Impaired in Aged Rats. J. Neurosci. 28, 7883–7890.

43. Middleton, S.J., Kneller, E.M., Chen, S., Ogiwara, I., Montal, M., Yamakawa, K., and McHugh, T.J. (2018). Altered hippocampal replay is associated with memory impairment in mice heterozygous for the Scn2a gene. Nat. Neurosci. 21, 996–1003.

## References

1. Skaggs, W.E., McNaughton, B.L., Gothard, K.M., and Markus, E.J. (1993). An information-theoretic approach to deciphering the hippocampal code. Adv Neural Inf Process Syst 5, 1030–1037.

2. O’Neill, J., Senior, T.J., Allen, K., Huxter, J.R., and Csicsvari, J. (2008). Reactivation of experience-dependent cell assembly patterns in the hippocampus. Nat. Neurosci. 11, 209–15.

3. Wills, T.J., Cacucci, F., Burgess, N., and O’Keefe, J. (2010). Development of the hippocampal cognitive map in preweanling rats. Science (80-.). 328, 1573–1576.

4. Davidson, T.J., Kloosterman, F., and Wilson, M.A. (2009). Hippocampal Replay of Extended Experience. Neuron 63, 497–507.

5. Silva, D., Feng, T., and Foster, D.J. (2015). Trajectory events across hippocampal place cells require previous experience. Nat. Neurosci. 18, 1772–1779.

6. Zhang, K., Ginzburg, I., McNaughton, B.L., and Sejnowski, T.J. (1998). Interpreting neuronal population activity by reconstruction: unified framework with application to hippocampal place cells. J.Neurophysiol. 79, 1017–1044.

7. Feng, T., Silva, D., and Foster, D.J. (2015). Dissociation between the Experience-Dependent Development of Hippocampal Theta Sequences and Single-Trial Phase Precession. J. Neurosci. 35, 4890–4902.

8. Drieu, C., Todorova, R., and Zugaro, M. (2018). Nested sequences of hippocampal assemblies during behavior support subsequent sleep replay. Science (80-.). 362, 675–679.

9. Zar, J.H. (2010). Biostatistical Analysis 5th ed. (Prentice Hall).

